# REALM: AO-based localization microscopy deep in complex tissue

**DOI:** 10.1101/2020.06.12.147884

**Authors:** Marijn E. Siemons, Naomi A.K. Hanemaaijer, Maarten H.P. Kole, Lukas C. Kapitein

## Abstract

Performing Single-Molecule Localization Microscopy (SMLM) in complex biological tissues, where sample-induced aberrations hamper detection and localization, has remained a challenge. Here we establish REALM (Robust and Effective Adaptive Optics in Localization Microscopy), which corrects aberrations of ≤1 rad RMS using 297 frames of blinking molecules to improve single-molecule localization. We demonstrate this method by resolving the periodic cytoskeleton of the axon initial segment at 50 μm depth in brain tissue.

## Results and discussion

Single-Molecule Localization Microscopy (SMLM) [1-3] enables exploring the nanoscale organization of cellular structures through repetitive localization of different sparse subsets of fluorophores. While this has provided many new insights into the organization of individual cultured cells, performing SMLM deep inside tissue or organoids has remained challenging. Accurate localization requires sufficient signal to noise and a non-aberrated point-spread-function (PSF) [4], both of which are compromised when imaging in biological tissue, in which a range of distinct cellular components cause complex light scattering [5].

A way to overcome this is the use of intensity-based Adaptive Optics (AO), which uses a deformable mirror in the emission path to compensate for tissue-induced wave-front distortions. The required shape of the mirror can be found iteratively by optimizing the contrast of the image (or guide star when possible) [6]. However, in SMLM the acquisitions are noisy and contain a strongly fluctuating amount of signal photons, rendering traditional approaches unusable. Various AO-methods have been proposed to overcome these issues [7-9], but the exact performance of these different AO-methods has not been consistently established and it has remained unclear what level of aberrations these methods can correct and under which conditions. Here we introduce REALM (Robust and Effective Adaptive Optics in Localization Microscopy), which corrects aberrations of up to 1 rad RMS using 297 frames of blinking molecules, thereby enabling robust SMLM at 50 μm depth and even up to 80 μm depth when pre-correcting spherical aberration.

Aberrations alter the points spread function (PSF), which decreases contrast and the spatial frequency support (see Figure 1a-d). The key for intensity-based AO is to find a relevant metric, a quality measure computed from the acquisitions, in combination with an optimization algorithm to efficiently and robustly optimize the metric [10]. To reduce the effect of strongly fluctuating signal levels of the acquisitions, all existing methods propose a weighted sum of the Fourier transform of the acquisition as metric, while differing in the specific weighting of the spatial frequencies and in normalization (see insert Figure 1d). We termed these metrics M1, M2 and M3, corresponding to Burke et al. [7], Tehrani et al. [8], and Mlodzianoski et al. [9], respectively. Because SMLM acquisitions are comprised of pseudo-random point sources, these methods effectively probe and optimize the Magnitude Transfer Function (MTF). As optimization algorithm to maximize the value of the metrics, Burke et al. use model-based optimization, Tehrani et al. use particle-swarm optimization, and Mlodzianoski et al. downhill simplex optimization.

**Figure 1.**
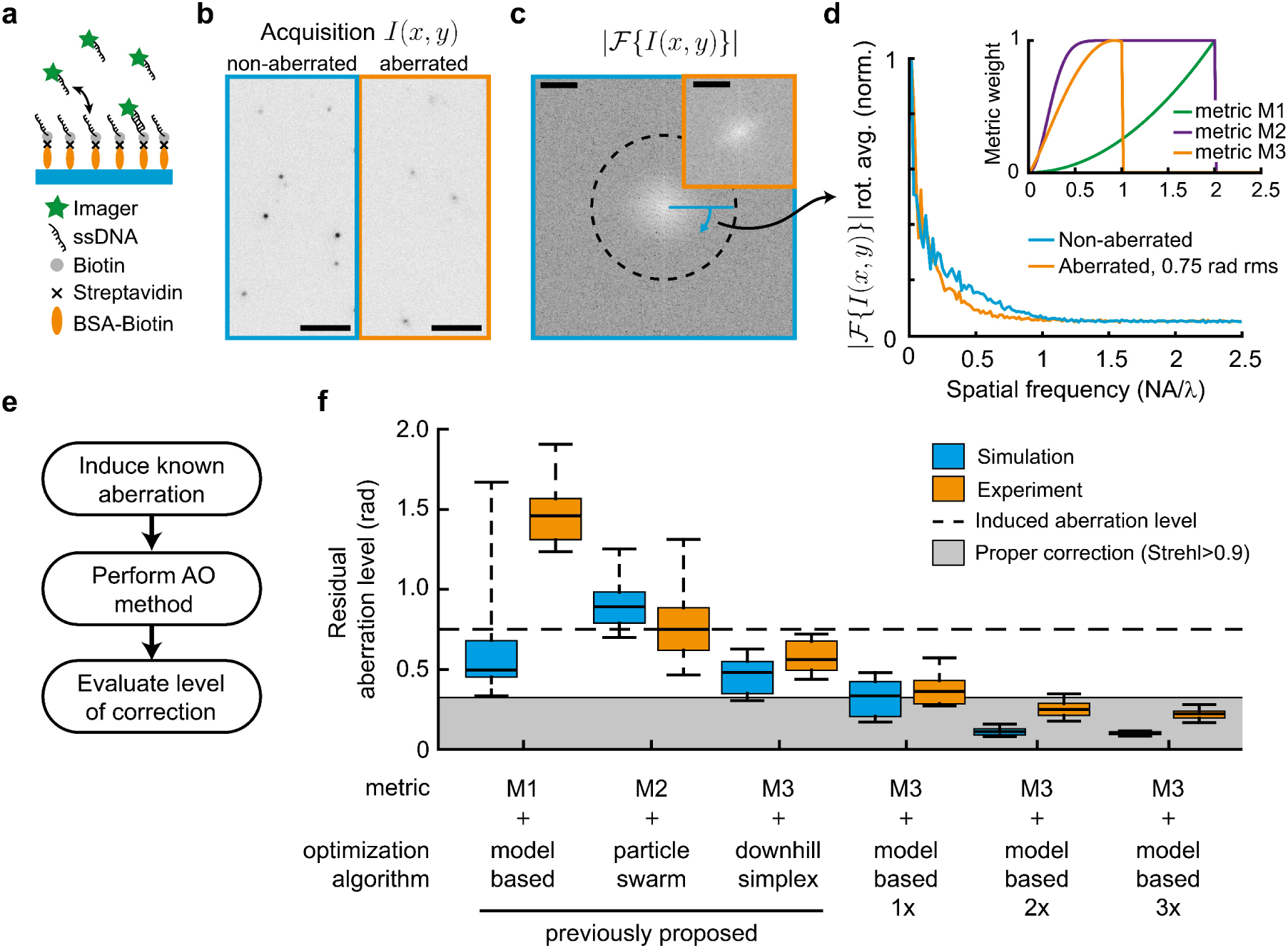
Systematic comparison between different AO methods reveals that only a new strategy achieves robust correction. **a**) DNA-PAINT test sample: Imager strands bind transiently to the coverslip, mimicking single-molecule blinking with consistent signal levels for many hours. **b**) Example acquisition of a non- and 0.75 rad RMS aberrated acquisition. Scale bar indicates 5 μm. **c**) 2D Fourier transform of a non-aberrated acquisition and aberrated acquisition (orange insert, center crop). Dashed line indicates 2NA/λ. Scale bar indicates NA/λ. **d**) The rotational average of (c) shows that noise dominates spatial frequencies above 1 NA/λ. The major decrease in the MTF due to aberrations occurs between 0.25 and 1 NA/λ. Insert shows the spatial frequency weights of the different proposed metrics. **e**) Strategy for comparing AO methods by inducing known aberrations in a well-corrected system. **f**) Performance of different AO metrics and optimization algorithms. Metric 3 in combination with model-based optimization leads to robust correction in 3 correction rounds (297 acquisitions). Boxplot indicates 9/91-percentile, 25/75-percentile and median for 25 random aberration configurations of 0.75 rad rms wave-front error consisting of random combinations of Zernike modes up to the fourth radial order (excluding piston, tip, tilt and defocus). Each frame (400×400 pixels, 26×26 μm) contains on average 13 emitters, emitting 2500 photons with a background of 20 photons per pixel.

To facilitate a systematic comparison between different methods, we used a DNA-PAINT based sample where molecules transiently bind to the coverslip, mimicking blinking [11] (see Figure 1a). This sample maintains stable signal and noise levels during the complete experimental sequence of several hours (see Methods and Supplementary Figure 1). Using a setup with a carefully calibrated deformable mirror (see Methods and Supplementary Figures 2&3) we introduced aberrations of known amplitude and assessed how well the various methods were able to correct these aberrations (see Figure 1e,f). We induced 25 random aberration configurations of 0.75 rad RMS wave-front error, consisting of random combinations of Zernike modes up to the fourth radial order (excluding piston, tip, tilt and defocus) and examined how well the different methods could estimate and correct the wave-front distortion. In addition, we tested the performance of the methods on simulated data sets (see Methods).

In both approaches, we found that the previously proposed methods were unable to meaningfully correct the aberrated wave-front. For example, method 1 and method 2 increased the aberration level in 100% and 48% of the experimental cases, respectively, whereas method 3 decreased the aberration level only by 20% (0.16 rad RMS) on average. The experimentally achieved corrections deviated to some extent from the simulation results (Figure 1f), likely due to additional noise sources such as camera read noise. Furthermore, the small initial aberration introduced by the DM limited the experimentally achievable aberration level to 0.2 rad RMS (Supplemental Figure 1e).

To understand why correction often fails, we next examined how the different metrics depend on noise levels and aberrations. The spatial frequency content of acquisitions in non-aberrated and aberrated conditions revealed that, in both cases, spatial frequencies above 1 NA/λ are dominated by noise (see Figure 1d). Metric M1 and M2 have the highest weights for these frequencies, which introduces a large amount of noise to the metric value. In contrast, M3 only weights the low spatial frequencies (<1 NA/λ), where noise levels are much lower. This makes metric M3 the most robust measure of the MTF and explains why M3 enables consistent correction, albeit without reaching diffraction-limited imaging (here taken as a Strehl-ratio of 0.9).

We wondered whether the limited correction obtained using metric M3 might be caused by the simplex optimization used in this approach. Simplex optimization is sensitive to a local noise in parameters’ space as it only compares two values per optimization parameter. Here, we suggest a model-based optimization that iteratively corrects Zernike modes by applying a sequence of biases for each Zernike mode to be corrected. The metric values for these series of acquisitions are then fitted to a curve (the model or so-called metric curve) to find the optimum (see Supplementary Figure 4) [12]. This procedure reduces noise and therefore appears more suitable for AO in SMLM. To test this, we implemented metric M3 in combination with model-based optimization. We first simulated aberrated acquisitions to assess the metric curve and found that, within a range of ±1 rad per Zernike mode, it could be approximated by a Gaussian function with offset (Supplementary Figure 5). Next, we used a series of simulations to optimize our method in terms of maximum bias range, the number of biases per Zernike mode, and the number of correction rounds (Supplementary Figure 6). This revealed that the precision with which a certain Zernike mode can be corrected depends on the amount of aberration in the other modes. Consequently, the use of multiple correction rounds improves the correction (see Figure 1f).

We experimentally verified this approach using the DNA-PAINT sample and were able to consistently reduce the induced wave-front error of 0.75 rad RMS by a factor of two in the first correction round (99 acquisitions). The second correction round (another 99 acquisitions) yielded further improvement and achieved diffraction-limited imaging in 21 out of 25 cases, whereas after the third correction round (297 acquisitions in total) all induced aberrations were corrected. Repeating this procedure for increasing amounts of induced aberrations revealed that this method is able to robustly correct aberrations of up to 1 rad RMS (see Supplementary Figure 7) in realistic signal and noise levels (2500 signal photons per emitter and 40 background photons). We termed this method REALM and implemented it as a freely available and open source Micro-Manager plugin (see Methods).

We next aimed to use this novel approach for deep imaging in complex brain tissue in which cell bodies, capillaries and fibres all may act as obstacles for light and distort the wavefront. To reduce the scattering induced by such samples we used a glycerol-based blinking buffer (see Supplementary Figure 8). We first performed SMLM on cultured COS-7 cells stained for microtubules. To mimic deep-tissue localization, we imaged through brain sections of 50 and 80 μm thickness (see Figure 2a-e). After performing REALM the state of the DM was alternated between the system-corrected state (without AO) and the sample-corrected state (with AO) every 500 frames (see Supplementary Movie 1 and Supplementary Figure 9). This revealed that with AO, the number of successful localizations was increased up to 6-fold, resulting in a strongly improved reconstruction. For the 80 μm thick slices, we estimated the spherical aberration to be around 100 nm and used this for precorrection. We switched the DM between this pre-corrected state and the sample-corrected state and again found an improvement (see Supplementary Figure 9), indicating that non-spherical aberrations substantially contribute to image deterioration.

**Figure 2.**
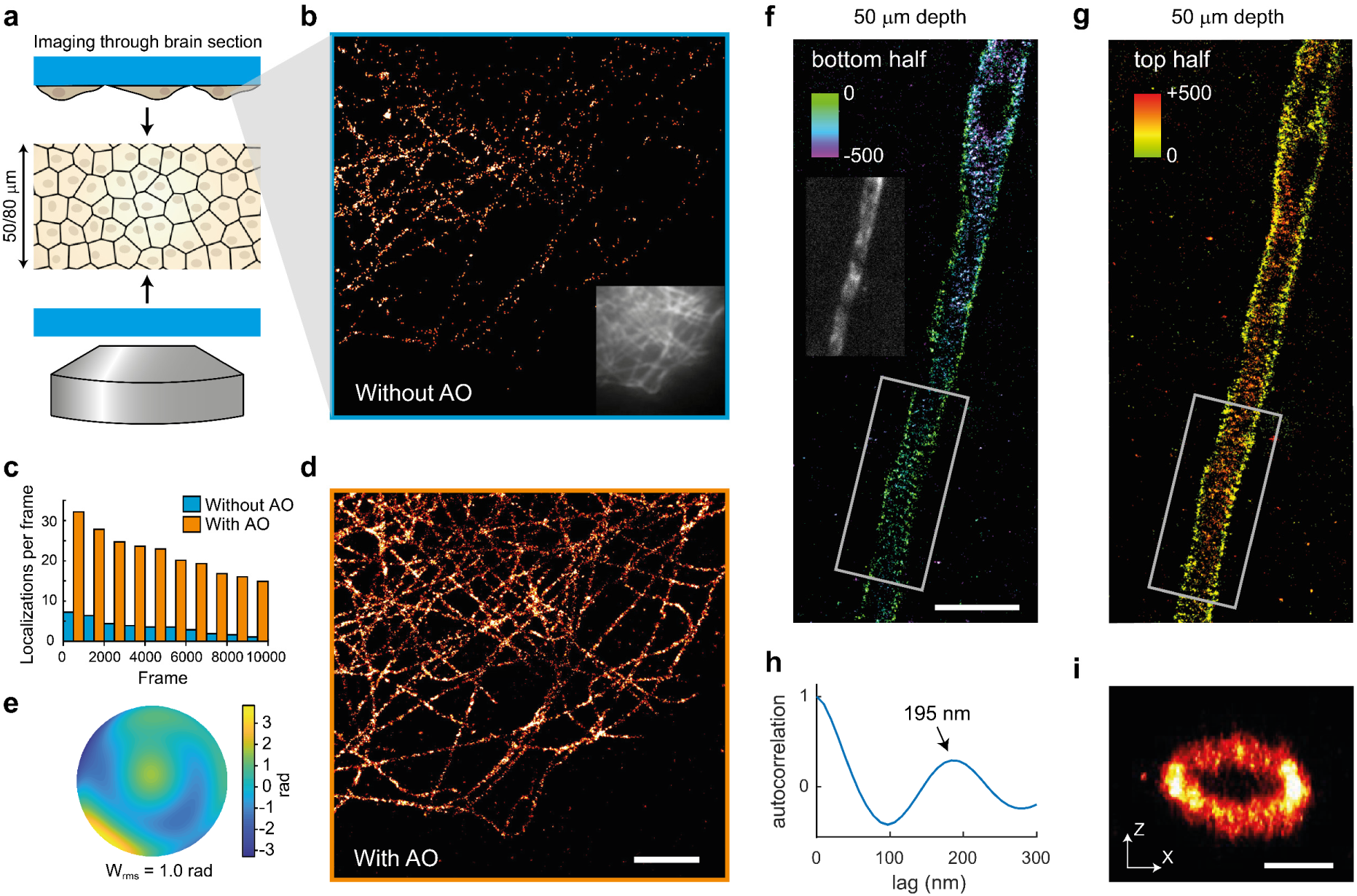
Improved single-molecule imaging through and in brain sections using adaptive optics. **a**) Illustration of the sample for panel (b-e). A 50 or 80 μm brain slice is mounted in between two cover glasses, on one of which COS-7 cells were grown. These cells were stained for microtubules and imaged through the brain section, to mimic deep-tissue imaging. **b**) SMLM reconstruction of microtubules in COS-7 cells through a 50 μm thick brain section with 5000 frames without AO sample correction. Insert shows the widefield image prior to correction. **c**) Number of localizations in the acquisition series during which the DM alternated between the system-corrected state (without AO) and sample-corrected state (with AO) every 500 frames. **d**) SMLM reconstruction of microtubules with sample-based correction for the same COS-7 cell as shown in (b). Scale bar indicates 2 μm. **e**) Estimated wave-front distortion as estimated using the AO correction method. **f**) SMLM reconstruction of the upper half of a layer 5 pyramidal neuron AIS stained for βIV-spectrin in a rat brain slice at 50 μm depth. Color encoding indicates the z-position. Scale bar indicates 2 μm. Insert shows the widefield image prior to aberration correction. **g**) Same as **f** but the lower half. The βIV-spectrin pattern are likely sites of synaptic connections. **h**) The average autocorrelation of the 12 line profiles shows a clear periodicity with the peak at 195±8 nm. **i**) z-x cross-section of the rectangular area indicated in **f**&**g**. Scale bar indicates 500 nm (both directions).

Finally, we aimed to image the axon initial segment (AIS) of layer 5 pyramidal neurons in rat cortical brain slices of 300 μm thickness. Landmark SMLM experiments have used neurons cultured on coverslips to reveal that axons display a ∼190 nm actin-spectrin based periodic structure called the membrane-associated periodic scaffold (MPS) [13], which at the AIS includes βIV-spectrin [13, 14]. However, due to the limiting imaging depth of conventional SMLM, linking these structures to functional assays of cells in their physiological anatomical connectivity (i.e. in brain slices) has remained challenging. In such slices, healthy neurons, as identified by electrophysiological experiments, are typically located at >30 µm depth from the slice surface and can be reliably targeted by a patch pipette up to 100 µm depth [15]. Using REALM, we performed multiplane 3D astigmatic SMLM imaging [9] on βIV-spectrin stained brain sections up to a depth of 50 μm, which enabled us to resolve the periodic patterning of this scaffolding protein in 3D and revealed a periodicity of 195±8 nm (mean±s.d., see Figure 2f,g,i, Supplementary movie 2&3 and Supplementary Figure 10). Importantly, we also successfully resolved the MPS of a functionally identified pyramidal neuron in a brain slice (see Supplementary Figure 11), demonstrating the feasibility of directly correlating functional studies to the nanoscopic architecture of the AIS.

In conclusion, we demonstrated that REALM can robustly correct aberrations in realistic signal and noise levels up to a depth of 50 μm in tissue. Therefore, our approach complements the recently introduced INSPR localization method, which demonstrated accurate 3D localization in aberrated conditions when imaging below 20 μm depth [16]. We anticipate REALM can enable 80 μm deep imaging in tissue when combined with approaches that limit out-of-focus fluorescence, e.g. selective plane illumination [17] or sparse labeling using an in vivo knock-in approach [18].

## 1. Methods

### 1.1 Set up

Experiments were performed using a Nikon Ti Eclipse body with a 100X 1.49 NA objective and a quad-band filter cube (containing a ZT405/488/561/640rpc dichroic and ZET405/488/561/640m emission filter, Chroma), to which a MICAO adaptive optics module containing a MIRAO52E (Imagine Optics) deformable mirror was mounted [19] (see Supplementary Figure 2). Detection is performed with a CMOS camera (Orca Flash v4.0, Hamamatsu). For excitation we used a single mode 647nm laser (140mW, LuxX, Omicron) and 405nm laser (60mW LuxX, Omicron) which can be used for normal widefield and TIRF illumination.

### 1.2 Calibration of the DM

In order to ensure accurate modulation of the Zernike modes, the deformable mirror needs to properly calibrated. The DM was calibrated by replacing the camera with a Shack-Hartmann sensor (HASO, Imagine Optics) to directly measure the wave-front, using the provided software. A 1 μm bead (TetraSpeck, ThermoFisher, T7282, dilution 1:1000) dried on a coverslip and mounted in glycerol was used as a point source. Mounting in glycerol reduces apodization in the pupil plane due to possible super-critical angle fluorescence, ensuring a homogeneously filled pupil. We blocked other beads in the FOV by placing an iris in the intermediate image plane. The wave-front deformation of each actuator was measured with 10 push-pull cycles to obtain the interaction matrix, which was then converted into the Zernike-based control matrix.

The calibration was verified using a phase retrieval algorithm [20] in combination with through-focus scans of 175 nm green fluorescent beads (PS-speck, ThermoFisher, P7220, dilution 1:500) mounted in PBS. This revealed that upon startup the shape of the mirror is imperfect due to thermal drift and needs to be corrected (see Supplementary Figure 3). The inverse estimated Zernike coefficients from the phase retrieval algorithm were subsequently applied by the mirror, after which a second through focus scan was acquired. This revealed that this approach was able to correct the complete system within 0.1-0.2 rad RMS (15-30 mλ) wave-front error. Prior to each experiment we performed this calibration step to ensure that the system was properly corrected. Next we modulated individual Zernike modes and acquired through focus scans of these PSFs. Phase retrieval indicated that all modes up to the 4^th^ order could be accurately modulated with little crosstalk between Zernike modes (see Supplementary Figure 3).

### 1.3 DNA-PAINT sample and imaging protocol

Sample chambers were prepared using double sided tape to create two cavities of 10 μl between a microscope slide and a #1.5 plasma-cleaned high-precision cover glass. Next, 10 μl of a solution of BSA-biotin (1 mg/ml in ultra-pure water MQ, Sigma Aldrich, A8549) was incubated for 5 min after being washed with 50 μl washing buffer (1x PBS containing 10 mM MgCl_2_). 10 μl of streptavidin (1 mg/ml in MQ, Sigma Aldrich, 434302) was then flushed in and incubated for 5 minutes and washed with washing buffer. Next biotin conjugated to the complementary DNA-strand P1 (1 mg/ml in MQ) [11] was incubated for 5 minutes and washed away. Finally, washing buffer containing 500 pM of Atto645 conjugated to DNA-strand I1 and green fluorescent beads (PS-speck, ThermoFisher, dilution 1:1000) was flushed in, after which the cavity was sealed with grease and nail polish. The fluorescent beads were used to monitor the stability of the deformable mirror and if needed to correct for thermal drift in between experiments.

For each AO correction, a random aberration configuration consisting of Zernike modes Z2±2, Z3±1, Z3±3, Z40, Z4±2 and Z4±4 (11 modes) was induced by the DM. The amplitudes of these modes were chosen uniformly random and normalized to 0.75 RMS rad. Next the adaptive optics method was performed (see below for implementation details). Afterwards the residual aberration level *W*_rms_ was evaluated as

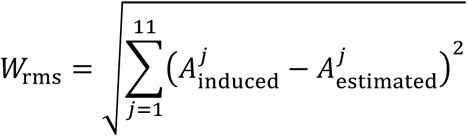

with 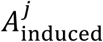 being the induced known Zernike coefficient of Zernike mode *j* and the 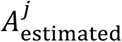 the coefficient of Zernike mode *j* estimated by the AO method. Prior and during the experiment the signal levels and state of the DM were monitored to ensure equal comparison between these methods (see Supplementary Figure 1 for experimental details).

### 1.4 Single-molecule acquisition simulations

The single-molecule acquisitions were simulated using a vector PSF model to capture the full complexity of the aberration configurations [21]. Blinking dynamics play a large role in the variability of the metric value and needed to be incorporated in the simulation. We mimicked these blinking dynamics by introducing a variability in the number of emitters in each frame and in the number of photons each emitter emits. The parameters for these distributions corresponded to the experimental signal levels of the DNA-PAINT sample (see Supplementary Figure 1). The number of emitters per frame was randomly chosen from a Poisson-distribution (with an average of 13 emitters), and the number of photons of each emitter followed an exponential distribution (with an average of 2500 photons). The emitters were randomly positioned with uniform probability across in the field of view (400×400 pixels, 65 nm pixel size) with a uniform background of 20 photons per pixel. Lastly Poisson noise was added to represent the shot-noise.

The performance of the AO methods was evaluated similar to the procedure described above. Known aberration configurations were induced, followed by AO-based corrections using the different approaches. For comparison with our DNA-PAINT experiment all emitters were placed in focus (z-position = 0 nm) and simulated with a 1.49 NA objective, while for the metric curve and optimization of our method (see Supplementary Figures 3&4) the emitters were uniformly random positioned in the z-direction between ±200 nm at a depth of 20 μm with a 1.35 NA silicon immersion objective with refractive index matching.

### 1.5 AO methods

Model-based optimization iteratively corrects Zernike modes by applying a sequence of biases of the Zernike mode that is to be corrected. The metric values of these acquisitions are computed and a Metric curve is fitted to find the optimum (see Supplementary Figure 4). Metric curve fitting was implemented by least-squares fitting with a Gaussian function with an offset (4 fit parameters). The width and center of the fitted Gaussian were constrained to prevent that occasional outliers in the metric values resulted in extreme fit values. The width was constrained to [0.4, 1] rad and the center to [-0.5, 0.5] rad. This greatly improved the performance of the previously proposed method [7] as the noise-sensitive metric M1 often led to extreme fit-values when not constrained. For Figure 1, the model-based optimization by Burke et al. was implemented with 11 biases per Zernike mode, whereas REALM used 9 biases per Zernike mode.

The downhill-simplex algorithm uses simplexes (higher dimensional triangles) to find an optimum in the parameter space. We implemented this optimization algorithm via the MATLAB function *fmincon* with the initial simplex size set to 0.2 rad, which was found to work optimally. Metric M3 was used for all Zernike modes as we did not induce secondary spherical aberration. Therefore, a separate simplex routine with a different metric for primary and secondary spherical aberration as originally proposed [9] could not be implemented. We did not find any reduced correction ability for primary spherical aberration using metric M3. However, we noticed that when correcting more than 4 Zernike modes, the simplex optimization was unable to converge in this larger noisy parameter space. Therefore, optimization was stopped after 300 acquisitions and the state with the best obtained metric value was taken as the estimated correction.

Particle-swarm optimization uses a collection of solutions moving through solution space, where their movement is affected by the individually best solution it found so far, as well as the groups best solution. Particle-swarm optimization was implemented via MATLAB *particleswarm* with a swarm size of 25 as suggested [8] with a maximum of 20 iterations for a maximum of 300 acquisitions. We used an initial swarm spansize of 0.1 rad and a maximum spansize of 0.75 rad. Other settings for *particleswarm* were set to standard values (*InertiaRange, SelfAdjustment* and *SocialAdjustment* set to 1).

### 1.6 COS7 staining

COS-7 cells for Figure 2a-e were seeded onto 25 mm coverslips. After 24 hours, cells were pre-extracted with 0.1% glutaraldehyde and 0.2% Triton-X100 in PEM80 (80 mM Pipes, 1 mM EGTA, 4 mM MgCl_2_, pH 6.8) for one minute. The cells were subsequently fixed with 4% PFA in PEM80 for 10 minutes. After washing in PBS (3×5 min) cells were permeabilized in 0.2% Triton-X100 in PEM80 for 15 minutes. After washing (3×5 min) blocking was performed in 3% BSA in PEM80 for 45 minutes and incubated overnight with a primary antibody against αTub (mouse IgG1, Sigma Aldrich, B-5-1-2, dilution 1:500). The cells were again washed with PBS (3×5 min) and incubated with secondary antibody (goat, anti-Mouse IgG (H+L), AlexaFluor647, Life Technologies, dilution 1:500) for 1 hour at RT. The coverslip was then placed on a microscope slide with the cells facing upwards, after which a 50 or 80 μm thick rat brain section (see below for details) was placed on the coverslip. The surplus of PBS was removed with a tissue and 70 μl of glycerol blinking buffer (see below) was deposited on the slice. Next a 25mm #1.5 high precision coverslip was placed on top of the slice and the assembly was sealed with nail polish. For the blinking buffer 10 μl of 1M MEA, together with 2,5 μl of 20% glucose and 1 μl of gloxy buffer (70 mg/ml glucose oxidase, 4 mg/ml catalase in MQ), were mixed with 86 μl of a mixture of 95% glycerol and 5% Tris 20mM, pH8.

### 1.7 Slice preparation and βIV-spectrin staining

All animal experiments were performed in compliance with the European Communities Council Directive 2010/63/EU effective from 1 January 2013. They were evaluated and approved by the national CCD authority (license AVD8010020172426) and by the Royal Netherlands Academy of Arts and Science (KNAW) animal welfare and ethical guidelines and protocols (IvD NIN 17.21.01 and 19.21.11).

To obtain sections with a fixed thickness (Figure 2a-e), rats were deeply anaesthetized by an i.p. injection of pentobarbital (50 mg/kg) and transcardially perfused with PBS and 4% PFA. The brains were removed and post-fixed in PFA for 24 hours after which the tissue was stored in PBS. Coronal sections of 50 μm thick were cut on a vibrotome (VT1000S, Leica Microsystems).

For βIV-spectrin staining (Figure 2f-i), rats were deeply anaesthetized by 3% isoflurane inhalation and decapitated, after which the brains were moved to ice-cold artificial cerebral spinal fluid containing (in mM): 125 NaCl, 3 KCl, 25 glucose, 25 NaHCO_3_, 1.25 Na_2_H_2_PO_4_, 1 CaCl_2_, 6 MgCl_2_, saturated with 95% O_2_ and 5% CO_2_ (pH 7.4). 300 μm thick parasagittal brain sections containing the primary somatosensory cortex were cut on a vibrotome (1200S, Leica Microsystems). Following a recovery period at 35 °C for 35–45 minutes slices were stored at room temperature in the ACSF. For whole-cell filling with biocytin (Supplementary Figure 10), the slice was transferred to a customized upright microscope (BX51WI, Olympus Nederland BV). The microscope bath was perfused with oxygenated (95% O_2_, 5% CO_2_) ACSF consisting of (in mM): 125 NaCl, 3 KCl, 25 glucose, 25 NaHCO_3_, 1.25 Na_2_H_2_PO_4_, 2 CaCl_2_, and 1 MgCl_2_. Patch pipettes were pulled from borosilicate glass (Harvard Apparatus, Edenbridge, Kent, UK) pulled to an open tip of 3 – 6 MΩ resistance. The intracellular solution contained (in mM): 130 K-Gluconate, 10 KCl, 4 Mg-ATP, 0.3 Na_2_-GTP, 10 HEPES, 10 Na_2_-phosphocreatine and 5 mg ml^−1^ biocytin (pH 7.25 adjusted with KOH, 280 mOsmol kg^−1^). An Axopatch 200B (Molecular Devices) was used to obtain whole-cell configuration. The cell was left to fill for 30 min, during which the bridge balance was monitored and stayed below 15 mΩ. Slices were fixed in 4% PFA (20 minutes) and blocked with 5% NGS and 2% Triton (2 hours) before incubation with rabbit anti-βIV-spectrin antibody in blocking buffer (1:1000, 24 hours, gift from M. Engelhardt). Slices were washed (3×15 minutes), incubated with goat anti-rabbit Alexa 647 (1:500 or 1:1000, 2 hours, ThermoFisher) and in the case of biocytin filling, with Streptavidin Alexa-488 conjugate (1:500, Invitrogen) and washed again. During all steps, the slices were at room temperature and on a shaker. The slices were stored in PBS (4°C). Before imaging, slices were incubated for at least 15 minutes in 95% glycerol and 5% Tris 20mM after which they were mounted in with between a microscope slide and #1.5 high precision coverslip with a 120 μm spacer (Secure-Seal Spacer, Thermofisher, S24735) in the blinking buffer described above.

### 1.8 SMLM detection, localization and reconstruction

The detection, localization and reconstruction was performed with the ImageJ plugin DoM (Detection of Molecules) [22] (see https://github.com/ekatrukha/DoM_Utrecht). DoM detects single molecules events by convolving images with a combination of a Gaussian and Mexican hat kernel. Localization is performed by an unweighted nonlinear 2D Gaussian fit with Levenberg–Marquardt optimization. SMLM reconstructions were rendered by plotting each molecule as a 2D or 3D Gaussian with standard deviations in each dimension equal to the corresponding localization errors. For astigmatic 3D localization the *z*-position was estimated from the difference in *x*- and *y*-width of the spot, where we corrected for depth-induced loss of astigmatism [19]. Drift was corrected by 2D cross-correlation of intermediate reconstructions consisting of 500 or 1000 frames.

### 1.9 REALM

REALM (Robust and Effective Adaptive optics in Localization Microscopy) is a free open-source Micro-Manager plugin (github.com/MSiemons/REALM) where the method described in this work is implemented (see Supplementary Figure 12). It offers a compact and intuitive user interface suitable for non-experts. It currently supports two types of DMs: MIRAO52E (Image Optics) and DMH40-P01 (Thorlabs, see github.com/HohlbeinLab/Thorlabs_DM_Device_Adapter for the device adapter) and we encourage others to build device adapters for other DM manufacturers to interface with REALM. The Fourier transform needed for the metric evaluation is implemented via the 2D Fast Hartley Transform (FHT), where the image is padded with the mode value of the acquisition to a size of 2^n^x2^n^ (with *n* in integer), usually resulting in a size of 256×256 or 512×512. All aberration correction for Figure 2 is performed with REALM.

---

**Supplementary figure 1.**
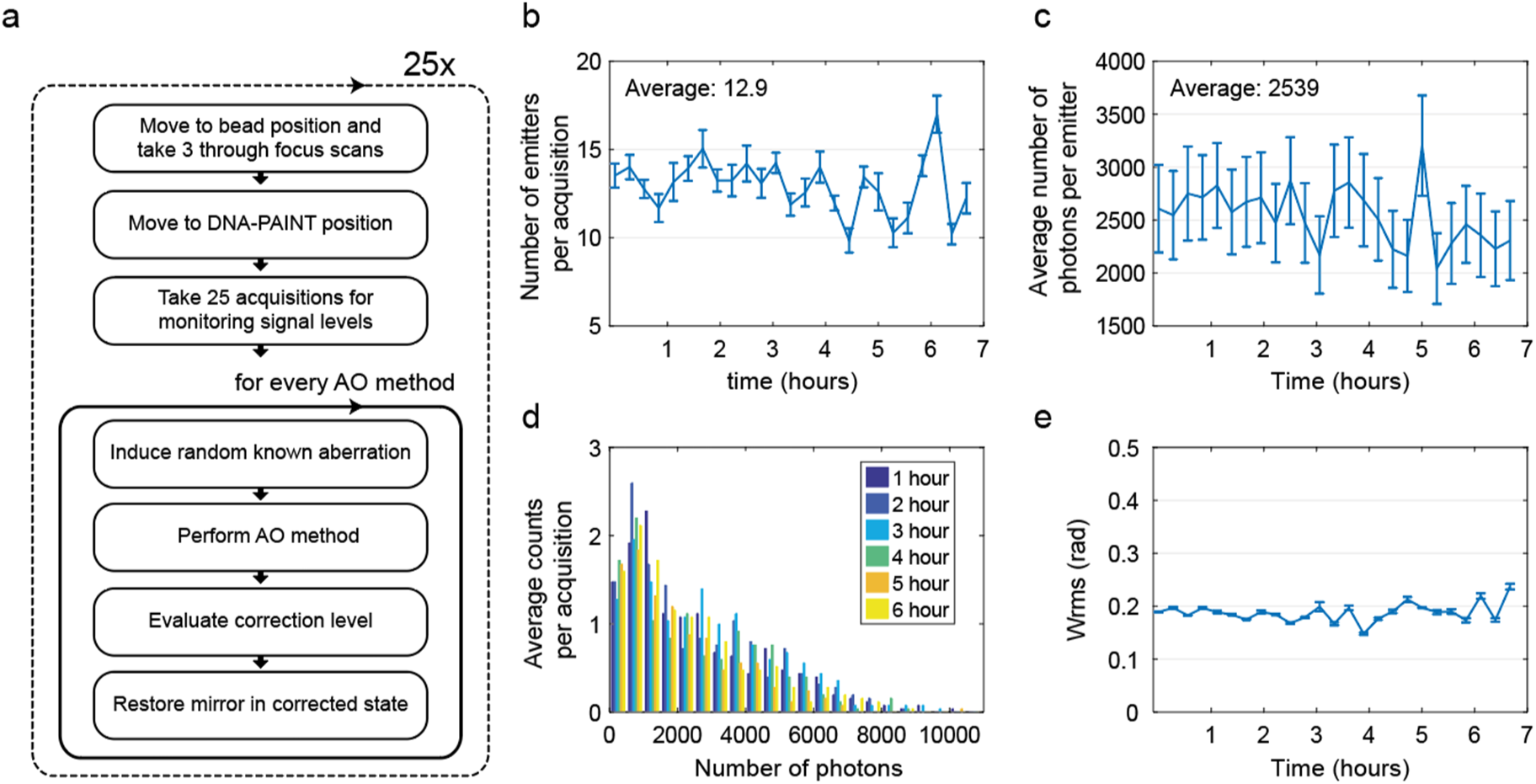
DNA-PAINT sample and deformable mirror remain stable during the experiment of figure 1f. a) Schematic of the experimental procedure, consisting of a monitoring part (first 3 blocks) and a correction part (bottom 4 blocks). For all AO methods tested, the stage moves to a position with a bead and acquires 3 through focus scan. Next, 25 frames were acquired at the DNA-PAINT position (without a bead in the FOV) with the system-corrected DM state. From these acquisitions the number of emitters (b), number of photons per emitter (c) and distribution of photon counts were measured (d). This revealed that the signal levels remained constant for the full 7 hour duration of the experiment. The acquired through-focus scans (in system corrected state) are analyzed with a phase retrieval algorithm [20] to check for possible drift in the mirror. The aberration level (e) remained at a level of 0.2 rad RMS during the whole experiment.

**Supplementary Figure 2.**
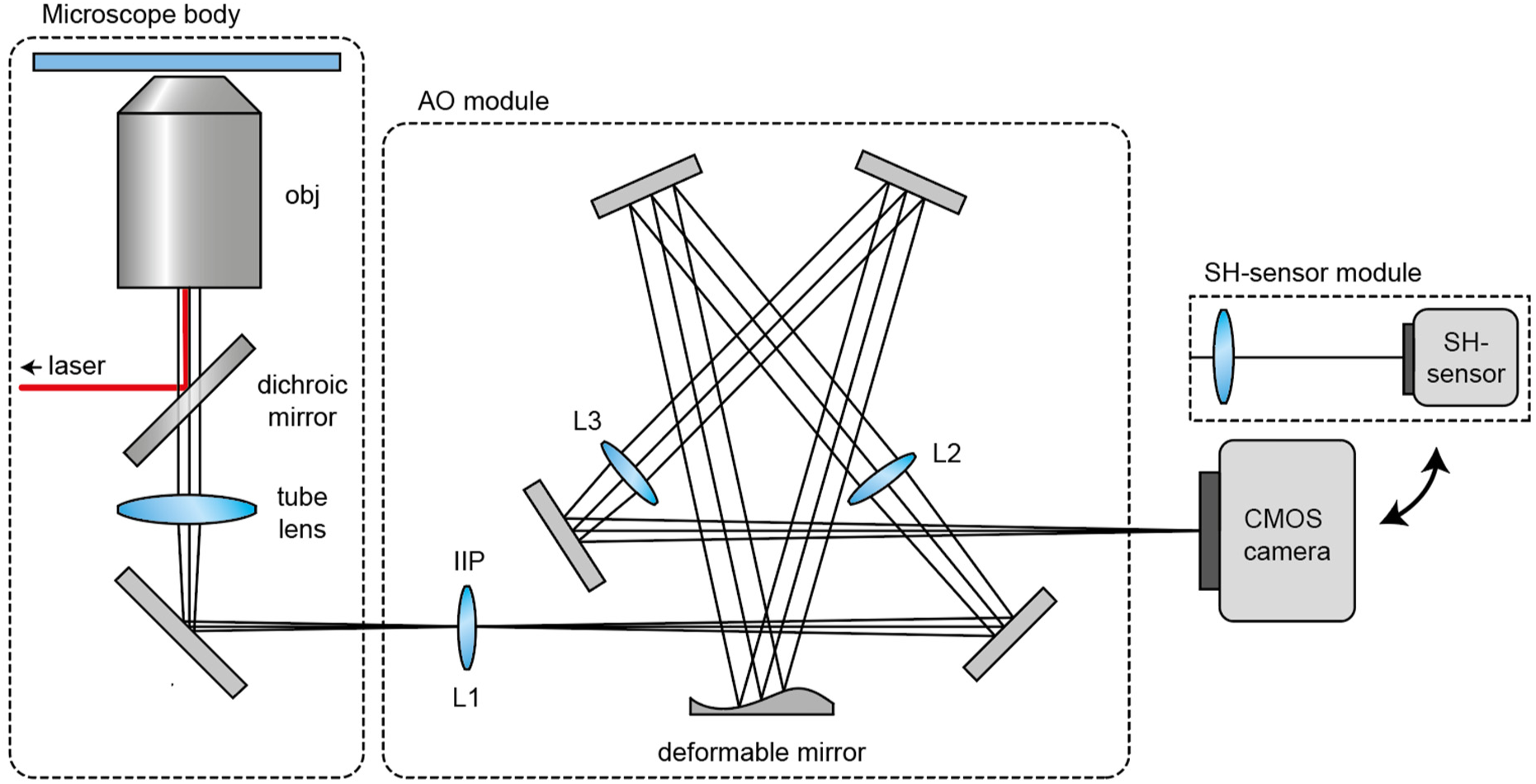
Illustration of the set up. The AO module consists of a 4F system (lenses L2 and L3, *f* = 500 mm), where the deformable mirror (DM) is placed in the back focal plane of L2. Another lens (L1, *f* = 750 mm) is placed in the intermediate image plane (IIP), which conjugates the pupil plane of the objective to the DM. This lens is needed as the tube lens and objective inside the microscope body are placed approximately 5 cm too close to each other to form a 4F system. For calibration of the DM, the CMOS camera is replaced with the SH-sensor module, which consists of a lens (*f* = 100 mm) and a Shack-Hartman sensor.

**Supplementary Figure 3.**
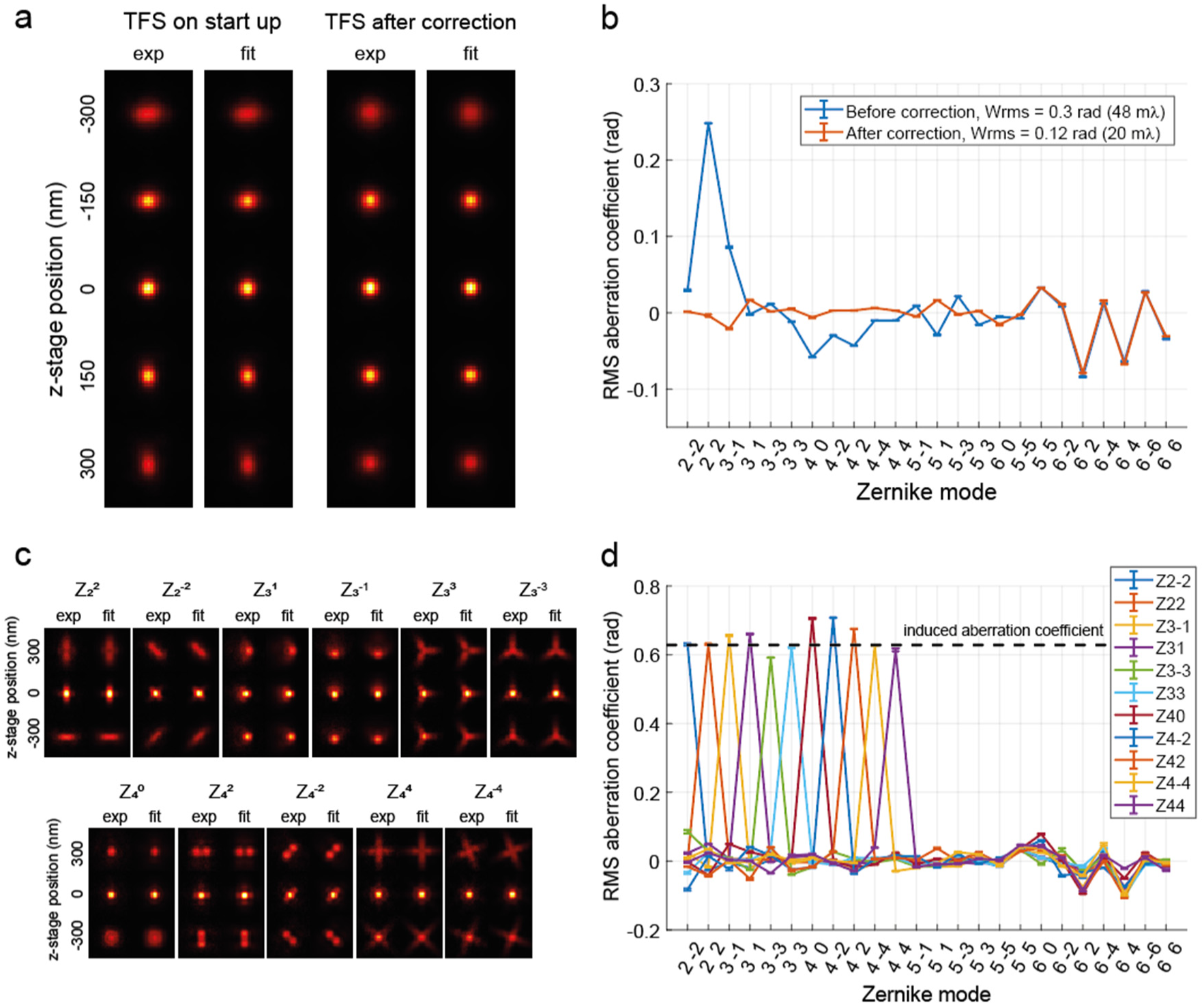
Verification of the DM-calibration. a) On start-up the experimental PSF (exp) was aberrated due to drift in the deformable mirror. A phase retrieval algorithm (fit) estimated the specific Zernike modes as shown in (b). The inverse Zernike coefficients of modes Z2±2, Z3±1, Z3±3, Z40, Z4±2, Z4±4, Z5±1, Z5±3 and Z60 were subsequently applied the mirror, which improved the PSF (TFS after correction). Phase retrieval revealed that all major contributing Zernike modes were nullified. c) PSFs and phase retrieval fits corresponding to Zernike modes Z2±2, Z3±1, Z3±3, Z40, Z4±2, Z4±4 with an amplitude of 0.63 rad (100 mλ). d) phase retrieval of (c) revealed that this DM is capable of modulating these Zernike modes accurately with little crosstalk.

**Supplementary Figure 4.**
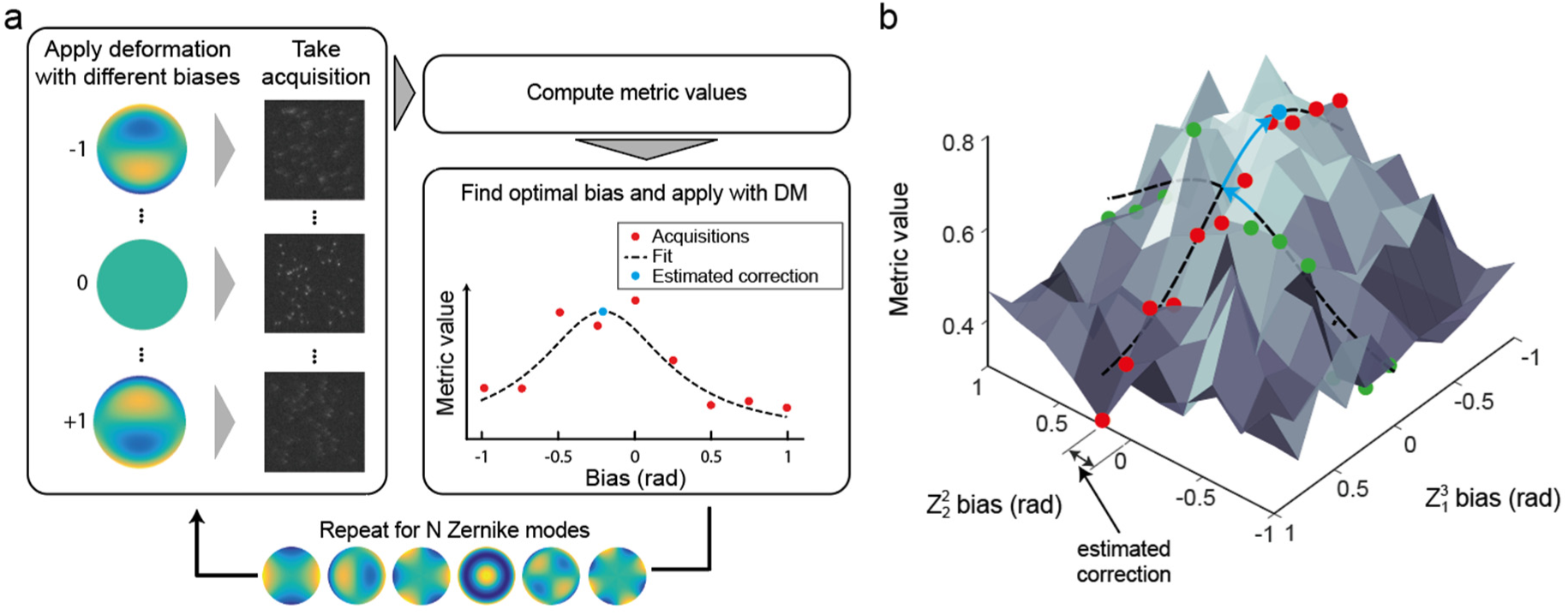
Schematic of model-based optimization. **a**) For each Zernike mode a series of frames is acquired with different biases (amplitude of the Zernike mode). From these acquisitions the metric value is computed. Next, the optimal bias is determined by fitting an appropriate function (the metric curve) to these points, after which the estimated correction is applied. This is performed for multiple Zernike modes. **b**) The optimum correction for Zernike modes can be more easily estimated when the contrast in metric value is improved. Therefore it is beneficial to first correct major expected types of aberrations, such as spherical aberration. For the same reason, multiple correction rounds also yield additional improvement.

**Supplementary Figure 5.**
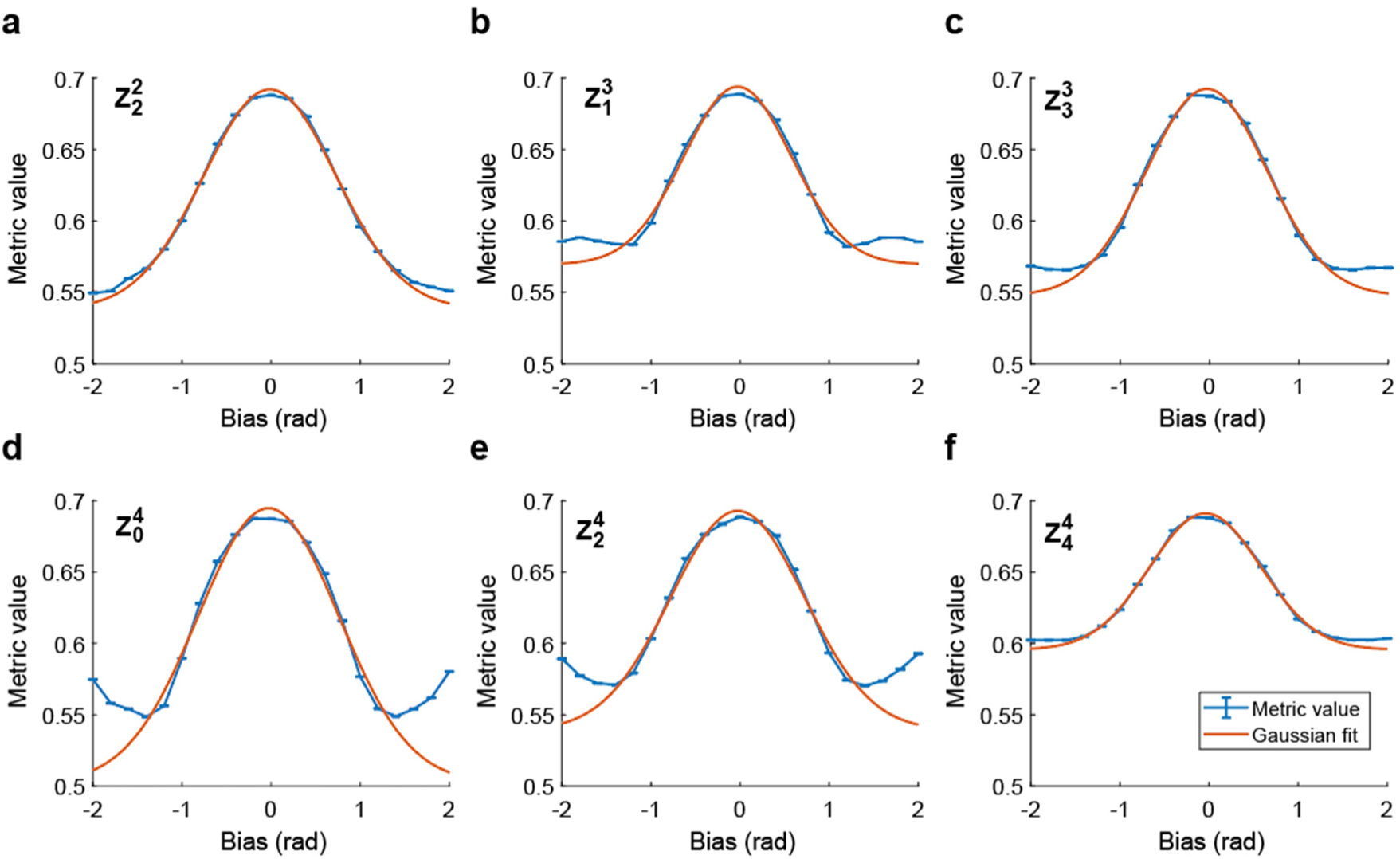
Metric curves for different Zernike modes. a-f) Metric value as function of applied bias for different Zernike modes. These values were obtained by simulating single-molecule acquisitions (see methods) with different biases for each Zernike mode. A Gaussian function with offset properly describes the metric values to all Zernike modes up to the 4^th^ order in a ±1 rad range and was therefore used as the metric curve. Increasing the bias beyond ±1.5 rad results in an increase in the metric value for some Zernike modes. This inversion of the metric value occurs due to contrast inversion at (specific) spatial frequencies.

**Supplementary Figure 6.**
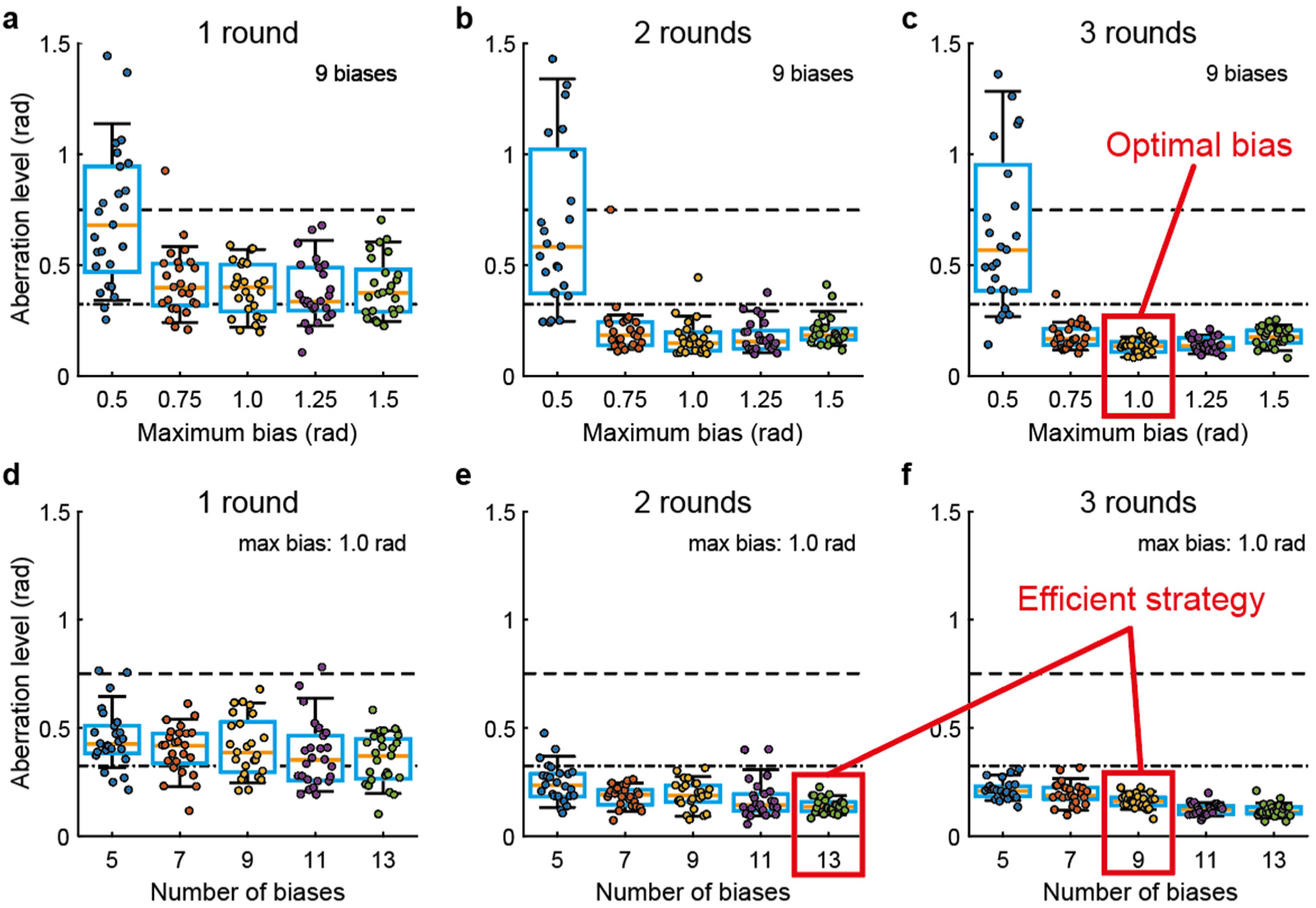
Simulation-based optimization of the model-based optimization algorithm. a-c) residual aberration level for different maximum applied biases and correction rounds, using 9 biases per Zernike mode (11 Zernike modes in total). Dashed line indicates induced aberration level (0.75 rad RMS). Residual aberrations are minimal when using a bias of 1 rad. d-f) Residual aberration level for different number of biases and correction rounds. Approaches with 13 biases in 2 correction rounds (total of 286 acquisitions for 11 Zernike modes) or 9 biases in 3 correction rounds (total 297 acquisitions for 11 Zernike modes) both constitute efficient strategies to achieve robust correction.

**Supplementary Figure 7.**
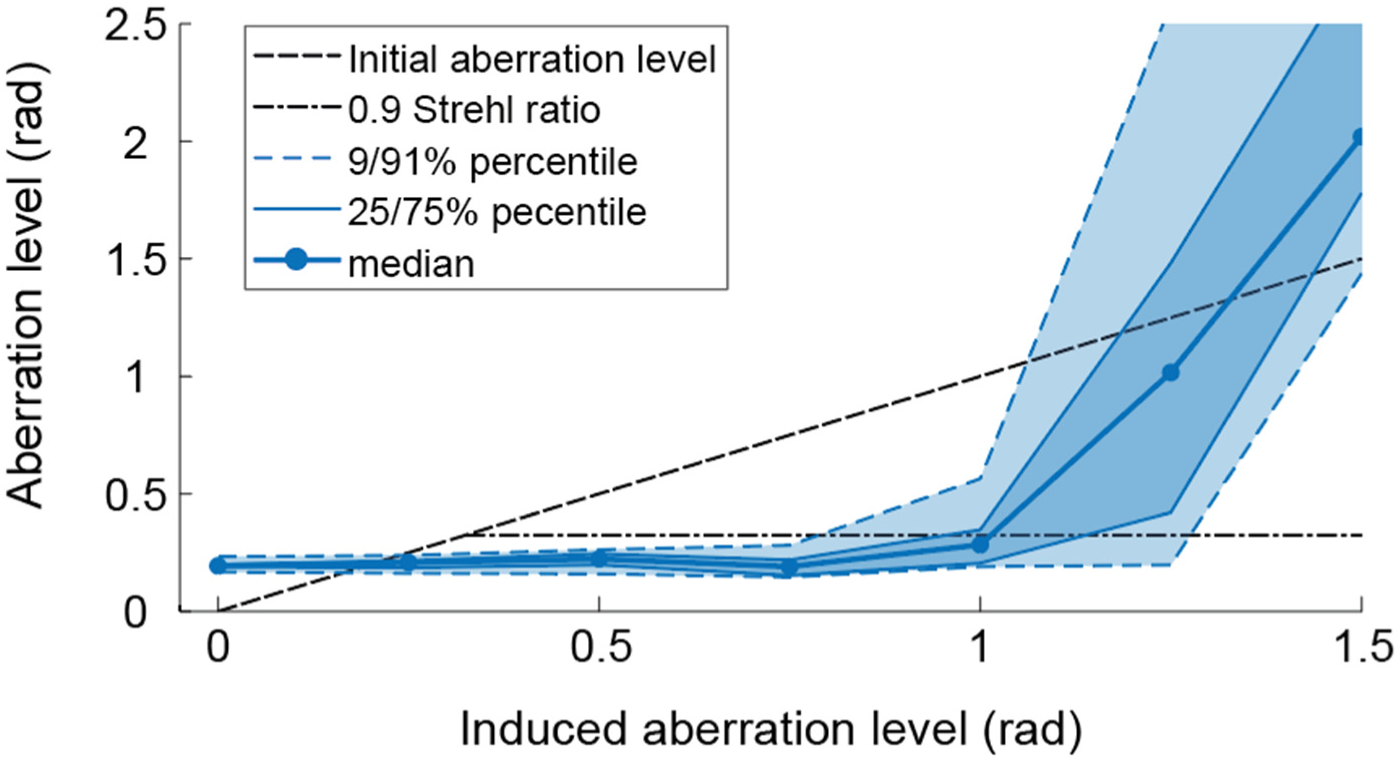
Experimental performance of REALM as function of induced aberration level using a DNA-PAINT sample. Here 2 correction rounds with 13 biases per Zernike mode with a maximum bias of 1 rad is used. For each aberration level, 20 random aberration configurations consisting of 11 Zernike modes (astigmatism, coma, spherical aberration, trefoil, second order astigmatism and quadrafoil) were induced by the deformable mirror and subsequently corrected. Results below the initial aberration level (dashed line) indicate improvement, results below 0.9 Strehl-ratio (dashed-dotted line) indicate proper imaging conditions. REALM was capable of correcting up to 1 RMS rad of wave-front error when the aberration was completely random and unknown. In practice, a major contribution is due to spherical aberration, which can be roughly pre-corrected, improving the performance. Signal levels for this experiment were 17 emitters per frame, emitting 2500 photons with a background of 40 photons per pixel.

**Supplementary Figure 8.**
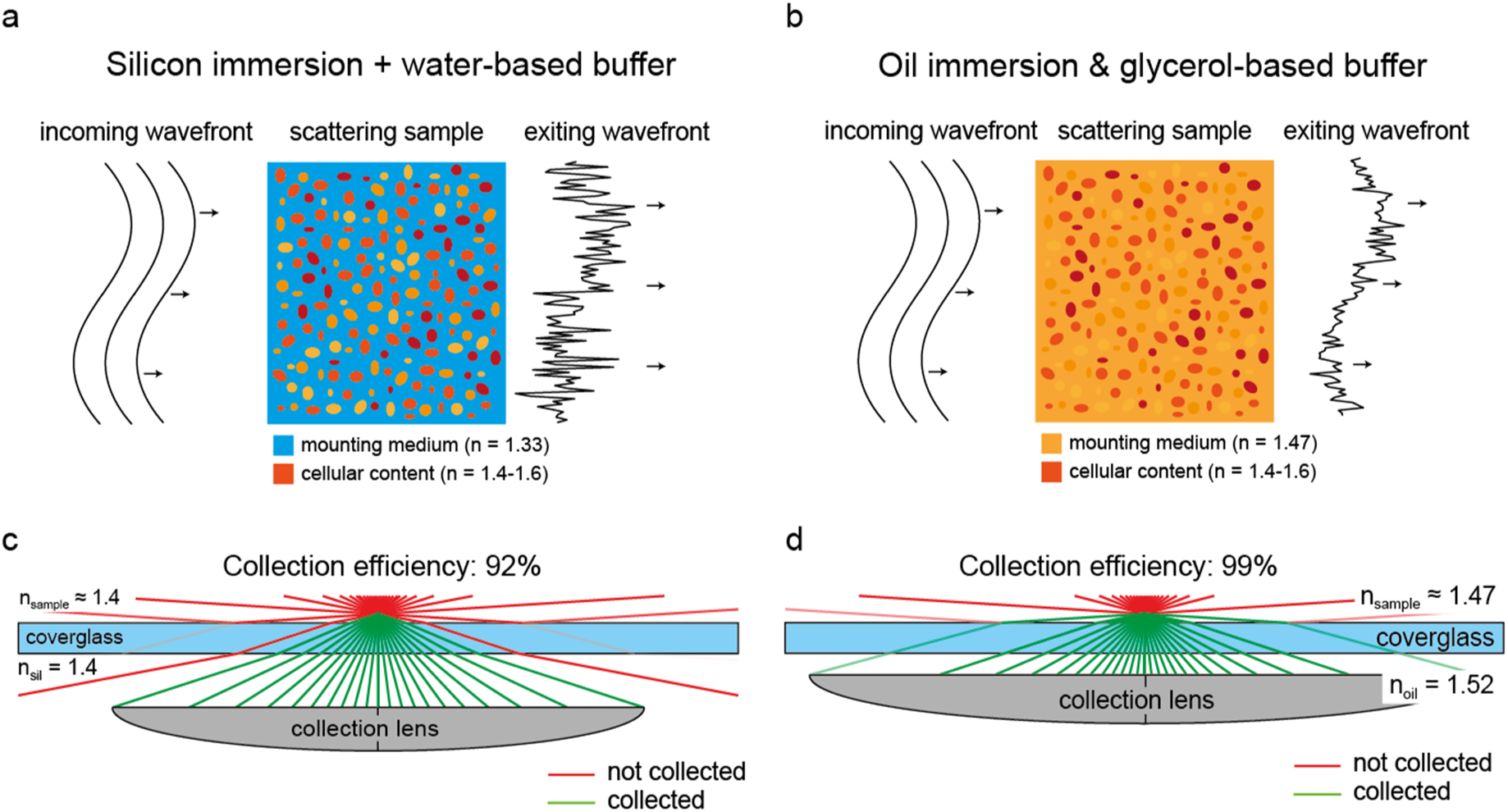
Illustration of the effect of a high refractive index buffer and objective choice. a) When using a water-based buffer the subcellular content of the tissue (organelles, DNA) has a large refractive index mismatch with the buffer. This local rapid change in refractive index drowns the gradually aberrated wave-front, thereby rendering AO less useful. b) By mounting the sample in a higher refractive index buffer using glycerol, the subcellular content causes less scattering and AO becomes more useful. c) A silicon immersion lens is the objective of choice when using a water buffer as the average refractive index of (brain) tissue is around 1.4. This matches the refractive index of the silicon oil, minimizing sample induced spherical aberration. However, as largest available NA (1.35) is smaller than the refractive index, the collection efficiency is only 92%. d) The glycerol-based buffer increases the average to around 1.47 (assuming a water content of 70%, which is replaced by the glycerol blinking buffer). Therefore a 1.49 NA oil immersion lens has a smaller refractive index mismatch than silicon oil and a higher collection efficiency as it collects the complete 2π sr solid angle and is therefore the objective of choice. For the computation of the collection efficiency we incorporated the Fresnel reflection at each interface.

**Supplementary Figure 9.**
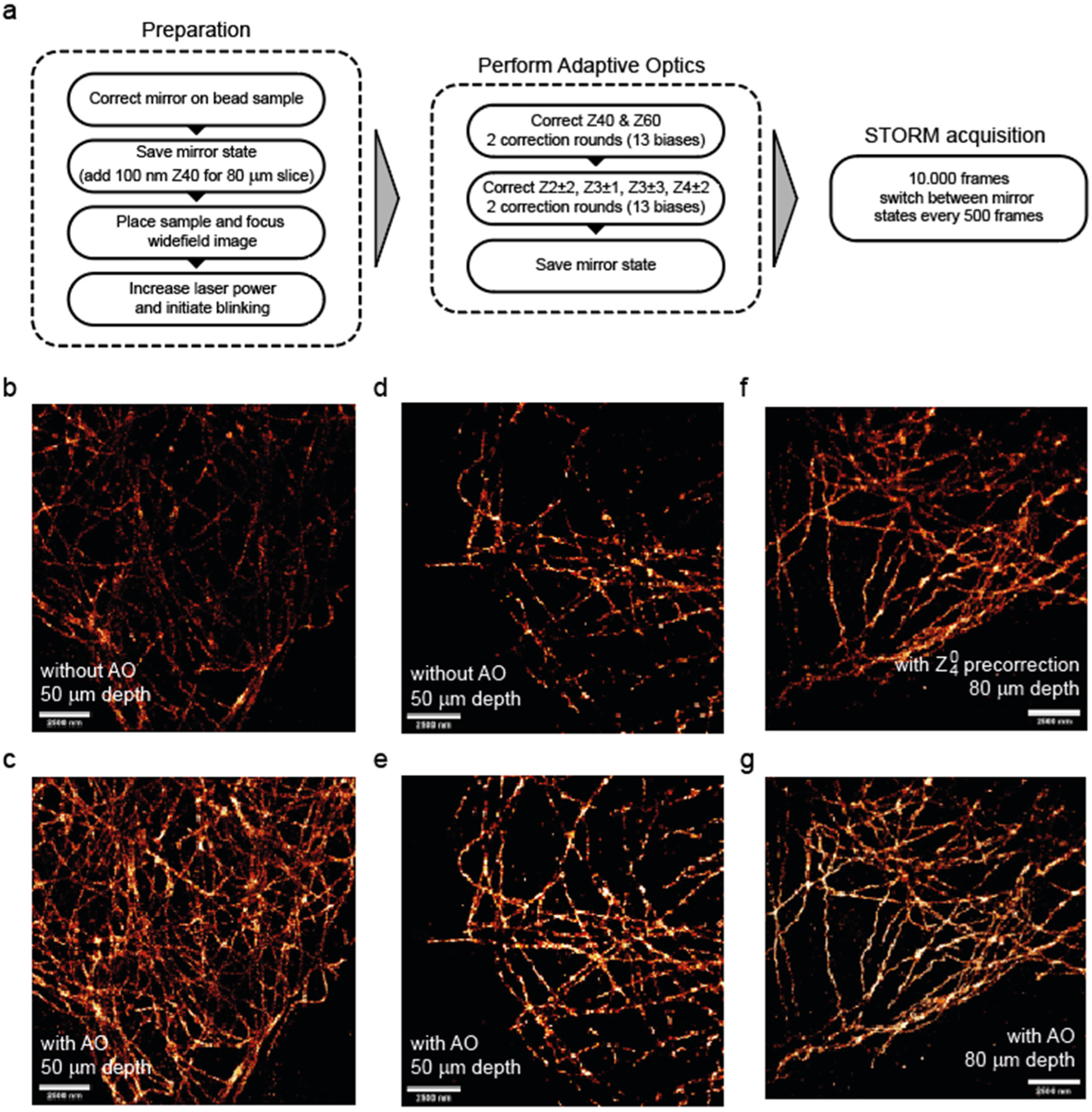
a) Experimental procedure for comparing SMLM with and without adaptive optics. First, the mirror was used to correct all system induced aberrations using a bead sample (see Supplementary Figure 2). This mirror state was saved and later used as the system-corrected DM state. For the 80 μm slice we pre-corrected spherical aberration by applying this mode with an amplitude of 100 nm (0.9 rad) RMS. Next the sample was mounted and focused using widefield imaging, while trying to keep the illumination minimal. Next the laser power was increased to initiate the blinking. Next, spherical aberration was corrected, followed by the other Zernike modes. After correction the mirror state was saved as sample-corrected state and the SMLM acquisition was started. During this acquisition the mirror switches between the system- or Z40 pre-corrected and sample-corrected state every 500 frames. b) SMLM reconstruction of microtubules in COS-7 cells imaged through a 50 μm thick brain section using the frames with the DM in system-corrected state (without AO). c) SMLM reconstruction of b) using frames with sample-corrected DM state. d&e) Another example as b&c. f) SMLM reconstruction of microtubules in COS-7 cells imaged through a 80 μm thick brain section using the frames with a precorrection of spherical aberration. g) as f but with frames with the DM in sample corrected state.

**Supplementary Figure 10.**
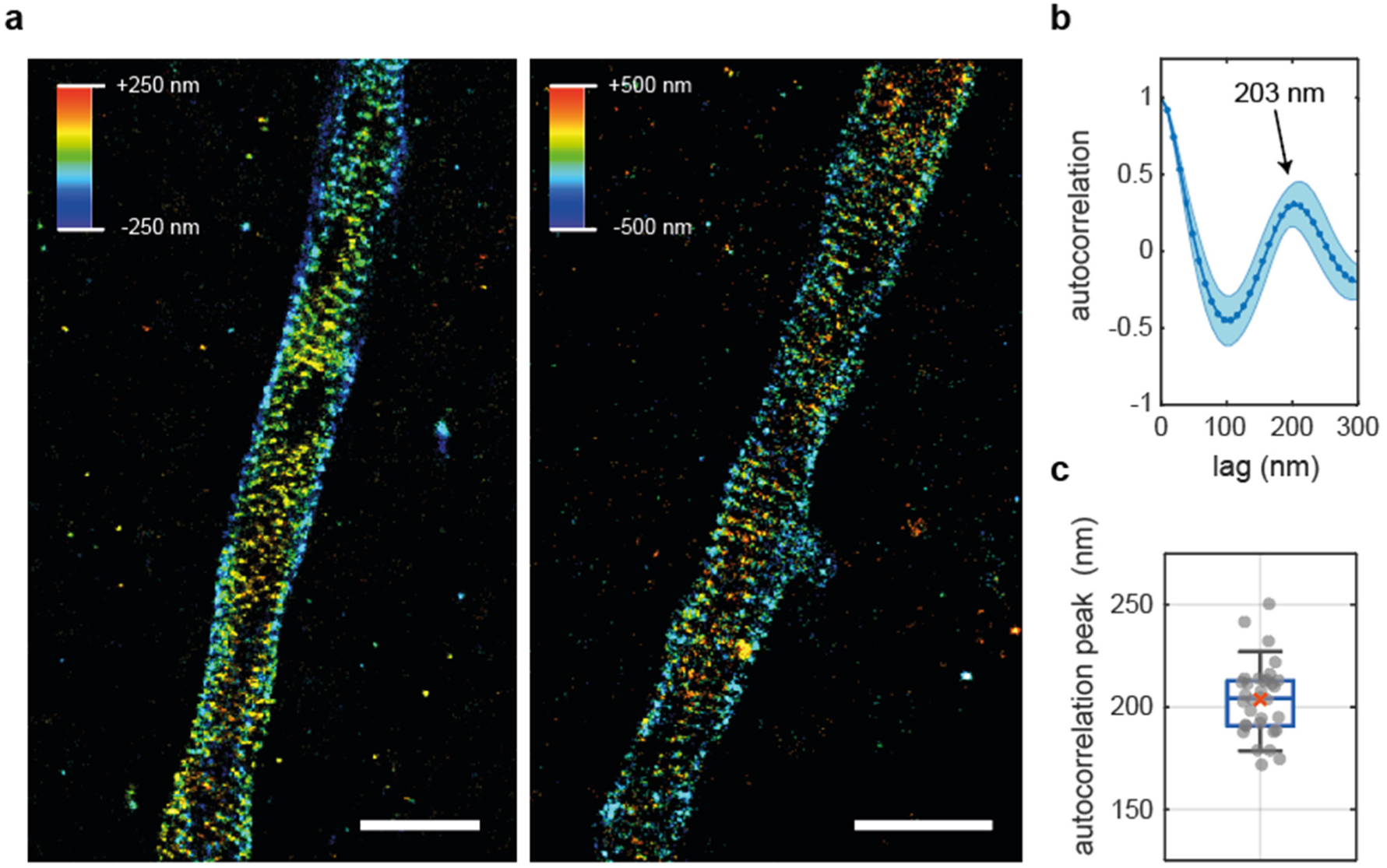
3D SMLM reconstruction of βIV-spectrin in the AIS. a) 2 SMLM reconstructions at a depth of 40 μm (left) and 50 μm (right). Scalebar indicates 2 μm. b) The average autocorrelation of 33 line segments in the reconstruction of (a) and Figure 2a&b shows a peak at 203±10 nm. Shaded region indicates the standard deviation. c) Distribution of the estimated autocorrelation peaks of all line segments. Boxplot indicates 9/91-precentile, 25/75-precentile and median,. Red cross indicates the mean.

**Supplementary Figure 11.**
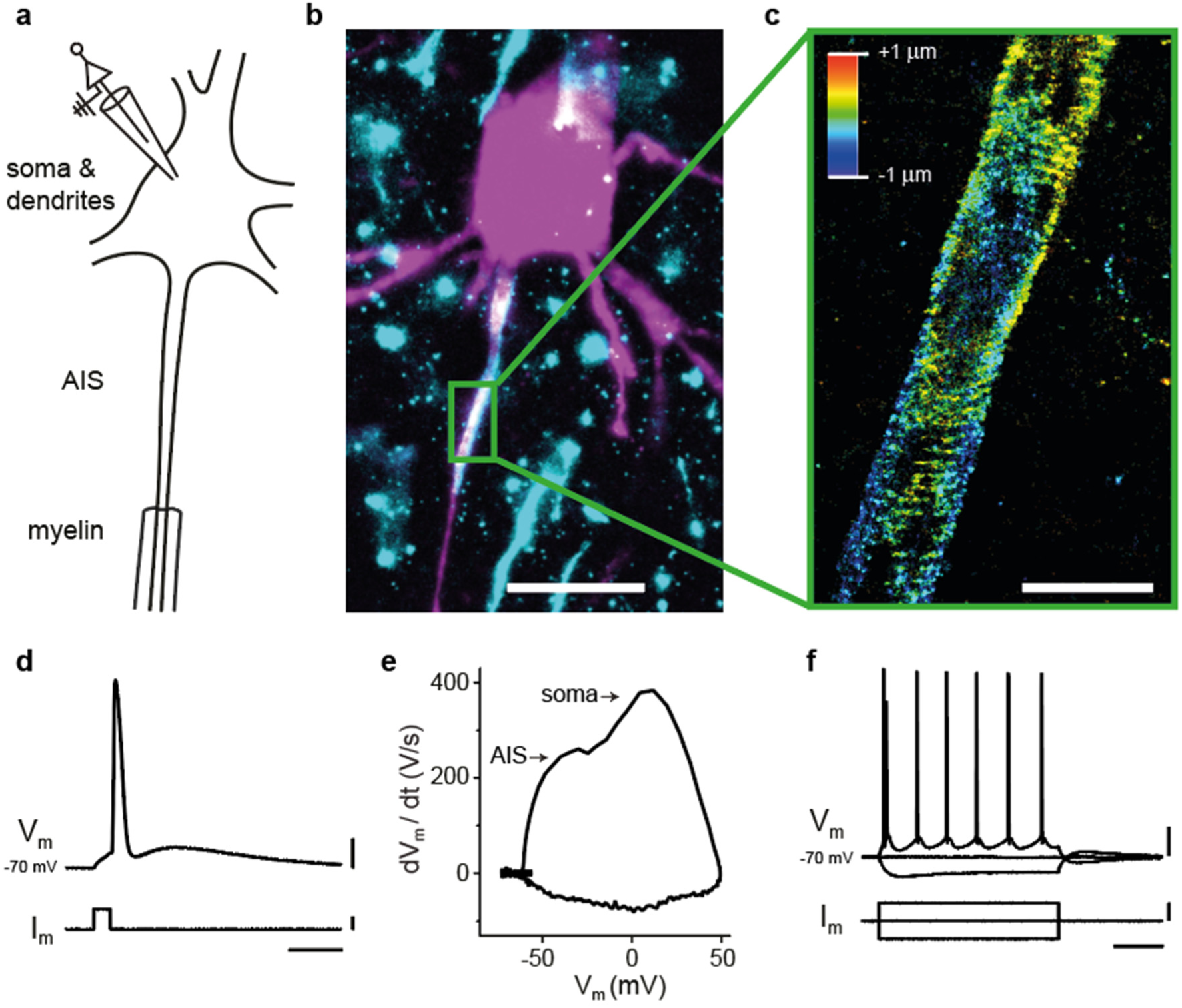
Combining 3D SMLM with electrophysiology. a) Schematic of whole-cell patch clamp recording of a layer 5 pyramidal neuron. b) Widefield image of a whole-cell recorded and biocytin-filled pyramidal neuron in a brain slice stained for βIV-spectrin. Magenta shows the cell morphology (streptavidin-Alexa488) and cyan shows the βIV-spectrin staining. c) 3D SMLM reconstruction of the βIV-spectrin at the AIS of the patched filled neuron. Here localizations of three focal planes are merged. Color indicates z-position. Scalebar indicates 2 μm. d) An action potential from the neuron presented in (b). V_m_ is the membrane potential and I_m_ the injected current, scale bars indicate from top to bottom: 20 mV, 500 pA and 10 ms. e) Phase-plane plot (dV_m_ /dt versus V_m_) of the action potential shown in (d) showing separate peaks from the AIS and the somatodendritic compartment. f) The membrane potential (V_m_) in response to negative and positive current injections (I_m_). Scale bars indicate from top to bottom: 20 mV, 200 pA, 200 ms.

**Supplementary Figure 12.**
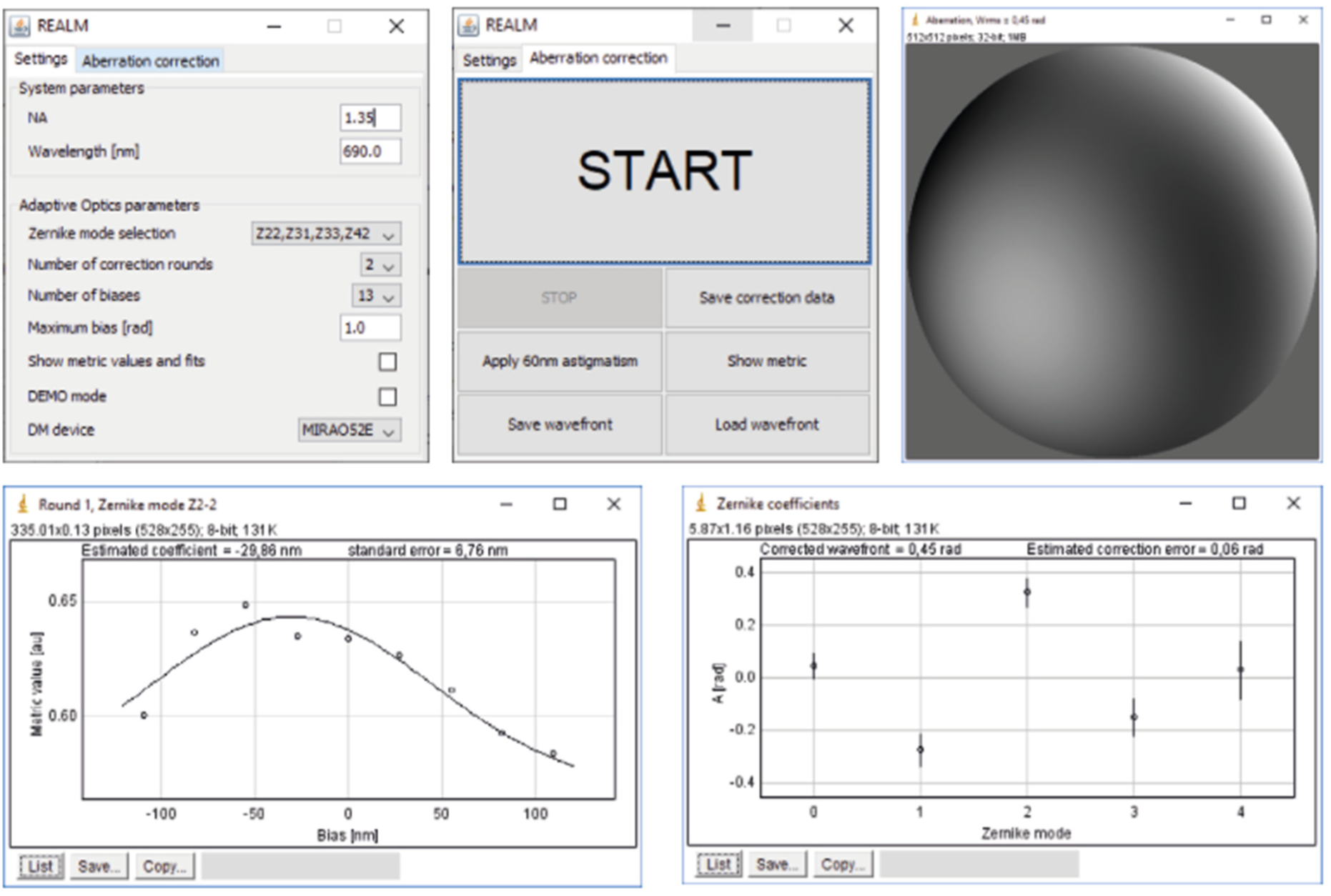
Images of the opensource Micro-Manager plugin REALM (https://github.com/MSiemons/REALM). Relevant parameters can be tuned (Zernike modes, number of biases, maximum bias, number of correction rounds). REALM requires only little input parameters (NA and wavelength), resulting in a clear and user-friendly interface.

