## Supplementary figures and images for "REALM: AO-based localization microscopy deep in complex tissue"

### Supplementary Movie 1

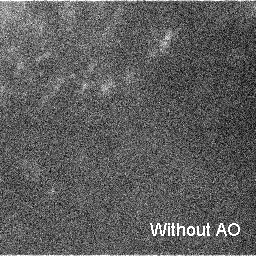

### Supplementary Movie 2

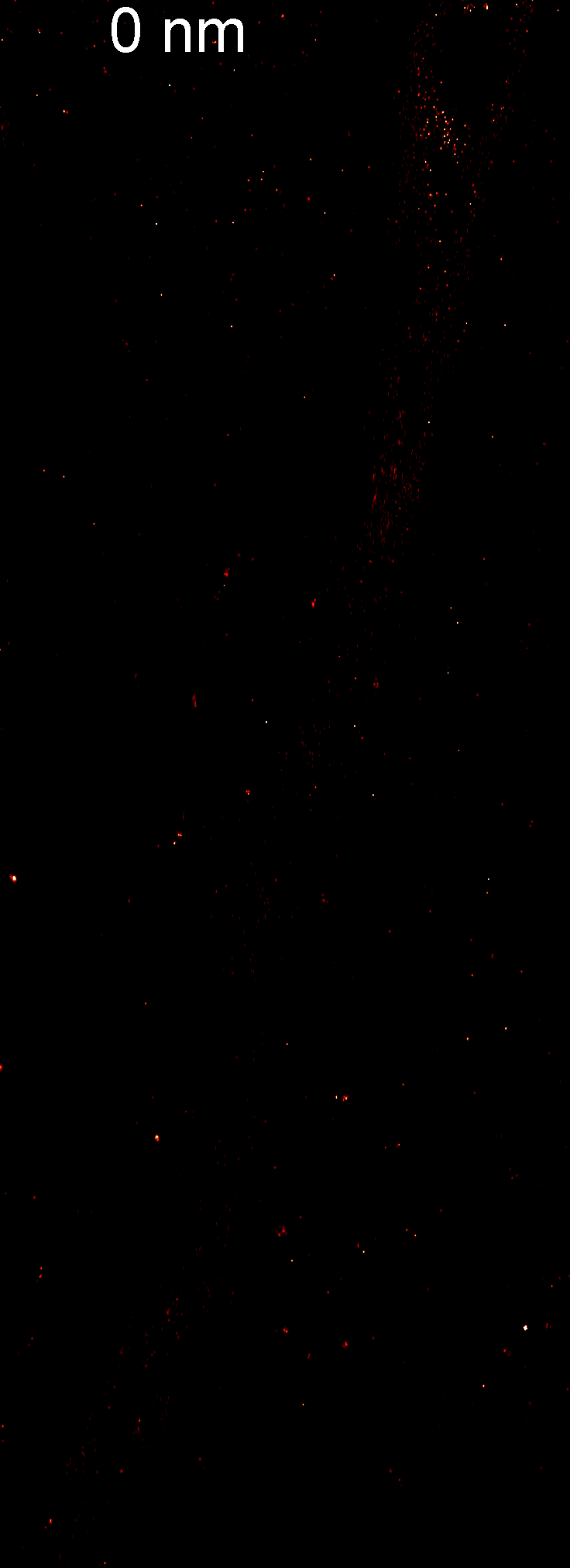

### Supplementary Movie 3

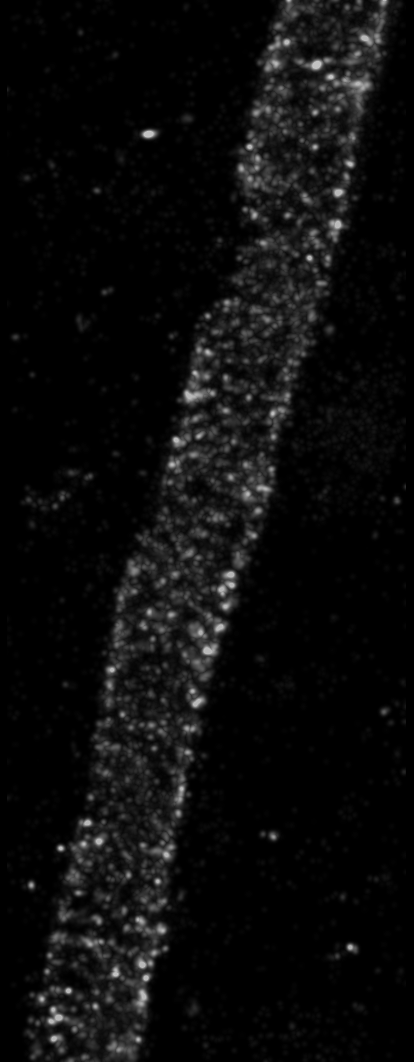
